# npphen: an R-package for non-parametric reconstruction of vegetation phenology and anomaly detection using remote sensing

**DOI:** 10.1101/301143

**Authors:** Sergio A. Estay, Roberto O. Chávez

## Abstract

For ecologists, the challenge at using remote sensing tools is to convert spectral data into ecologically relevant information like abundance, productivity or traits distribution. Among these features, plant phenology is one of the most used variables in any study applying remote sensing to plant ecology and it has formally considered as one of the Essential Biodiversity Variables. Currently, satellite imagery make possible cost-efficient monitoring of land surface phenology (LSP), but methods applicable to different ecosystems are not available. Here, we introduce the ‘npphen’ R-package developed for remote sensing LSP reconstruction and anomaly detection using non-parametric techniques. The package implements basic and high-level functions for manipulating vector and raster data to obtain high resolution spatial and temporal LSP reconstructions. Advantages of ‘npphen’ are: its flexibility to describe any LSP pattern (suitable for any ecosystem), it handles time series or raster stacks with and without gaps, and it provides confidence interval for the expected LSP at yearly basis, useful to judge anomaly magnitudes. We present two study cases to show how ‘npphen’ can successfully reconstruct and map LSP and anomalies for contrasting ecosystems.

## Introduction

The development of cost-efficient tools for monitoring the effects of climate change on vegetation and biodiversity has turned into a high priority in natural resources management, conservation and human welfare. In this scenario, remote sensing arises as a powerful tool for vegetation monitoring at large scales (Horning et al. 2010). Currently, dense time series of satellite imagery are freely available from missions such as Landsat or MODIS from NASA or the Sentinel from ESA allowing detailed description of natural and anthropogenic landscapes (Petorelli et al. 2014).

For ecologists, the challenge at using remote sensing tools is to convert spectral data into ecologically relevant information like abundance, productivity or traits distribution. Among these features, plant phenology is one of the most used variables in any study applying remote sensing to plant ecology and it has formally considered as one of the Essential Biodiversity Variables (EBV’s) for species traits’ monitoring (Pereira et al. 2013). An accurate reconstruction of the annual plant phenological cycle, valuable in itself, is also fundamental for detection and quantification of anomalies on primary production dynamics. From a remote sensing perspective, Land surface phenology (LSP) has been defined as the study of the spatio-temporal development of the vegetated land surface through the use of satellite sensors (de Beurs and Henebry 2005). Because anomaly detection requires the comparison with an expected pattern, LSP reconstructions should be the base of any study aimed at detecting changes in the spatial or temporal patterns of plant phenology (Hargrove et al. 2009, Verbesselt et al. 2012).

In this regard, Verbesselt et al. (2010, 2012) summarized the main challenges that LSP reconstructions and anomaly detection have to overcome. First, remote sensing observations are a combination of multiple signals acting at different conditions and time scales, and therefore, the ability of any method to detect a significant anomaly relies on its capacity to differentiate the expected phenological pattern from noise (aerosols, clouds, geometric errors, etc.). Second, change detection techniques should be independent of vegetation-specific thresholds. This means that anomaly detection should be based on the dynamics of the system itself, and not on predefined thresholds. Finally, the method needs to be robust to missing data, which is important for regions where climate prevents periodical records from optical sensors (e.g. at high latitudes).

Remote sensing based vegetation indices such as the Normalized Difference Vegetation Index (NDVI, Tucker 1979) or the Enhanced Vegetation Index (EVI, Huete et. al. 1994) provide multi-year pixel-base time series at high temporal and spatial resolution. LSP reconstructions using vegetation indices time series can be conducted using two approaches. First, by using a theoretical frequency distribution, which is considered an appropriate representation of the expected dynamic of the system. Usually, this is a function with an explicit seasonal component fitted to data using OLS. This theoretical pattern works by assuming that regular, annual waves are a good representation of the plant phenological cycle (e.g. Malo 2002, Verma et al. 2016), which is not true for all ecosystems. However, this approach is the most used in the literature (see de Beurs and Henebry 2010 for a review of LSP methods). The second approach, which we call empirical, uses the observed frequency values, and defines the expected distribution directly from observed data without reference to a theoretical model. The advantage of this approach is its flexibility to adapt to the particular conditions of every site, e.g. arid and semiarid ecosystems where seasonal approaches are not suitable (Beurs and Henebry 2010, Broich et al. 2015). At the best of our knowledge, this approach has never been used in LSP reconstruction.

Following this empirical approach, here we present a new computational methodology for LSP reconstruction and anomaly detection using remote sensing data, which takes advantage of the flexibility of kernel density estimation to overcome the challenges summarized by Verbesselt et al. (2010). We first introduce the mathematical basis of the method, followed by two examples of different plant phenology regimens and disturbances to demonstrate its performance. To facilitate the application of our method to users, we provide a complete implementation in the R environment: the ‘npphen 1.1-0’ package.

## Methods

### LSP reconstruction

For reconstructing LSP, the first step is to estimate the expected value of a vegetation index through the growing season. This in turn depends on the estimation of the probability density function *f*(x). In our method, we approximate *f*(x) by *ƒ̂*(x) using a Kernel Density Estimation (KDE) procedure.

Let us define X = (X_1_,…, X_n_), a sample time series containing paired values of a vegetation index (VgI), like NDVI or EVI, and time corresponding to the day of the growing season (ranging from 1 to 365, e.g. Julian day, Fig. 1b). So,

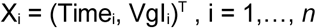

**Figure 1.**
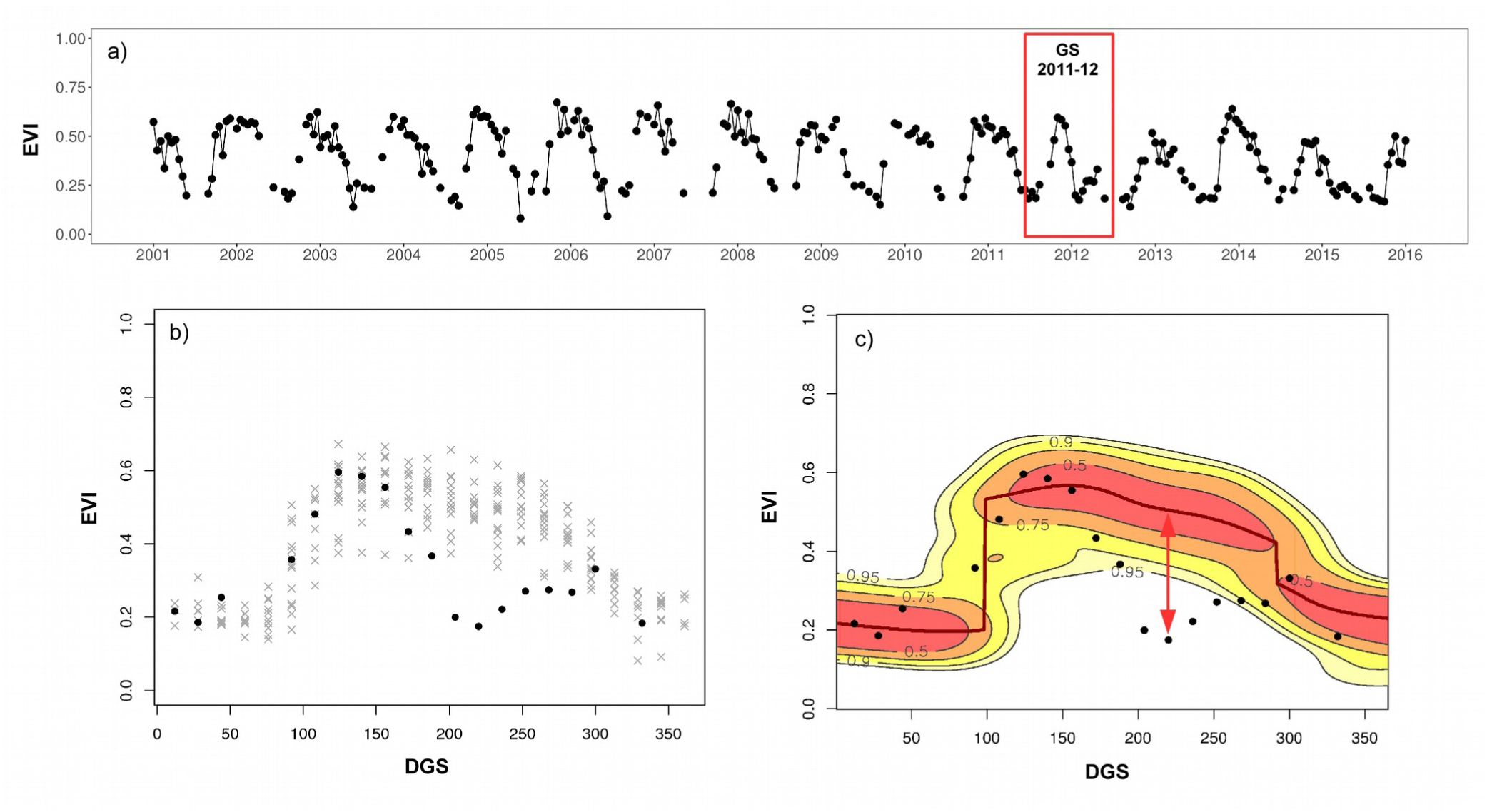
Example of npphen applied to time series of the Enhanced Vegetation Index (EVI) in the case study 1. a) Original time series of the Moderate Resolution Imaging Spectroradiometer (MODIS) 16-days EVI composites. The red box shows the growing season (GS) 2011-2012 when the outbreak of *Ormiscodes amphimone* took place. b) All EVI values from all growing seasons (grey x), except for the growing season to be analyzed (GS 2011-2012 in black circled dots), are used to reconstruct the leaf phenology of the deciduous *Nothofagus pumilio* forest. c) Frequency map of the EVI-time space displaying the probabilities of EVI values at different days of the growing season (DGS): the dark red line (maximum probability) is considered the reference EVI phenological curve. The difference between the dark red line (reference) and the black dots (observed values) are negative EVI anomalies (red arrow), which can be related to foliage loss due to the defoliating insect outbreak.

For example, if our dataset covers *p* annual phenological cycles with *m* observations per cycle, then our time series will contain *n*=*m*×*p* observations (Fig. 1a-b provides a time series example). However, for our method having the same number of points per cycle is not mandatory.

We define *ƒ̂*(x) as the bivariate density function of X estimated by

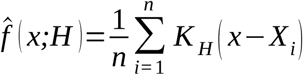

where x is a generic point in the bivariate Time-VgI space, X_i_ = (Time_i_, VgI_i_)^T^, H is the so called bandwidth 2×2 matrix, and K is the kernel. KDE in 2 dimensions works by centering a bivariate kernel (e.g. a Gaussian kernel) around each observation, and by averaging the heights of all kernels till obtaining the final density estimation. The size of the kernel in each dimension is defined by H. For more details about theoretical aspects of KDE see Wand and Jones (1994).

In our algorithm we used a Gaussian kernel over the observed VgI values. Estimation is performed over a grid of 365 columns (daily estimation) and 500 rows. The bandwidth is defined using the multivariate plug-in selector of Wand and Jones (1994). The bandwidth matrix defines the size of the kernel in each dimension (diagonal) and the rotation of the kernel in reference to the axes (anti-diagonal).

Using *ƒ̂*(x) we can identify the set of most probable values of VgI along the phenological cycle and its confidence interval (Fig. 1c). The set of VgI values representing the more probable value per day along the phenological cycle will be set as the expected LSP for this site. The anomaly is defined as the difference between the observed and the expected value at a given day (red arrow in Fig. 1c),

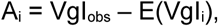

### Algorithm implementation

The algorithm has been implemented in the *Non-Parametric Phenology* “npphen 1.1-0” R-package (https://CRAN.R-project.org/package=npphen). The package implements basic and high-level functions for manipulating vector data (numerical series) and raster data (satellite derived products). Processing of very large raster files using multi-core computing functionalities is also supported. The package contains functions that reconstruct LSP from numerical series of vegetation index (e.g. Fig. 1a), but also from 3-dimensional stacks of images representing a vegetation index for a given area at a pixel basis. For an specific time period (e.g. 1 or 2 years), npphen also calculates the anomaly of the observed vegetation index given the reconstructed LSP (a description of the functions is provided in table 1).

**Table 1.** Brief description of the functions contained in the npphen package.

| **Functions** | **Description** |
| --- | --- |
| Phen | Reconstruct the annual LSP cycle from a time series of vegetation greenness |
| PhenAnoma | Calculate anomalies with respect to the regular phenological cycle using time series of vegetation greenness |
| PhenAnoMap | Calculate anomalies with respect to the regular phenological cycle using raster stacks of vegetation greenness |
| PhenKplot | Plot the most probable vegetation greenness values and their confidence intervals |
| PhenMap | Reconstruct annual LSP using raster stacks of vegetation greenness |

For functions implemented for numerical series (Phen and PhenAnoma) the output is a numerical vector containing the expected value of the vegetation index or the anomalies for each date during a given period (Fig. 1). Functions implemented for raster stacks (PhenMap and PhenAnoMap) generate as output a new raster stack with the same spatial dimensions than the input stack and a time dimension equal to the number of dates per growing seasons of the original data. Raster stack functions reconstruct LSP and calculate anomalies for each pixel independently.

The ‘**npphen’** R package, manual and examples of its use can be downloaded from The Comprehensive R Archive Network (https://cran.r-project.org/package=npphen).

## Comparison with other packages

Different from other R packages devoted to phenology, “npphen” reconstructs the expected “longterm phenology” and its confidence intervals for a given period of time (e.g. 34 years of NDVI GIMMS data), which can be used to set a robust base line and make comparisons to observed phenology at certain times (anomaly detection) or to analyze long-term changes in phenology (e.g. comparing the phenology of subsequent five-years periods). Other r packages such as “phenology” (Girondot 2018) are intended to interpolate or fit a parametric phenological curve to data for a single growing season or consecutive growing seasons (Girondot 2018). The “phenopix” package (Filippa et al. 2016), devoted to analyze digital pictures of vegetation cover, also includes some functions to fit parametric functions to the resulting phenological data. Others such as “bfast” (Verbesselt et al. 2010) or “green-brown” (Forkel et al. 2013) provide several tools to study land surface phenology using remote sensing time series data, but using a different approach as the one used here. Using parametric functions to fit the complete time series, they compute phenological metrics (e.g. start, end, peak and length of the growing season) per growing season, or in the case of “bfast”, decompose the series into a phenological (seasonal) component, a trend component, and a remainder error. Abrupt changes in trends can be related to vegetation perturbations, defining periods (between perturbations) for which the phenological signal is homogeneous and accounted by the parametric function. This approach does not deal well with large inter-annual variability in the time series, since no parametric function can describe subtle changes in the annual phenological cycle (Forkel et al. 2013).

## Study cases

### Temperate forests and insect outbreaks

We applied our algorithm to reconstruct the LSP of *Nothofagus pumilio* forests in Chile and to quantify the defoliation (negative anomaly) caused by outbreaks of the moth *Ormiscodes amphimone*. We used the 16-day compositions of MODIS EVI (Version 5), which is sensitive to vegetation green biomass (Huete et al. 2002). We downloaded 365 EVI images obtained from the Terra satellite (MOD13Q1 product) covering the region of Aysen, Southern Chile. All scenes were pre-processed considering the quality assessment bands and using the ‘raster’ package (Hijmans and Van Etten 2014) in the R software (R Core team 2017). From this stack of satellite imagery, the expected LSP cycle of the forest was reconstructed per pixel. An example of the LSP cycle and its confidence interval for an specific pixel is shown in Fig. 2d. The reconstructed phenological curve resembles the seasonal behavior of this deciduous species. Furthermore, an outbreak of *O*. *amphimone* reported on January 2012, was detected by means of EVI anomalies (Fig. 2d). By applying the anomaly mapping function ‘PhenAnoMap’ from the npphen package we obtained a 23-layer stack (one layer per date from the MODIS data) containing anomaly values per pixel. EVI anomaly maps showing defoliation levels are shown in Fig. 2a-c.

**Figure 2.**
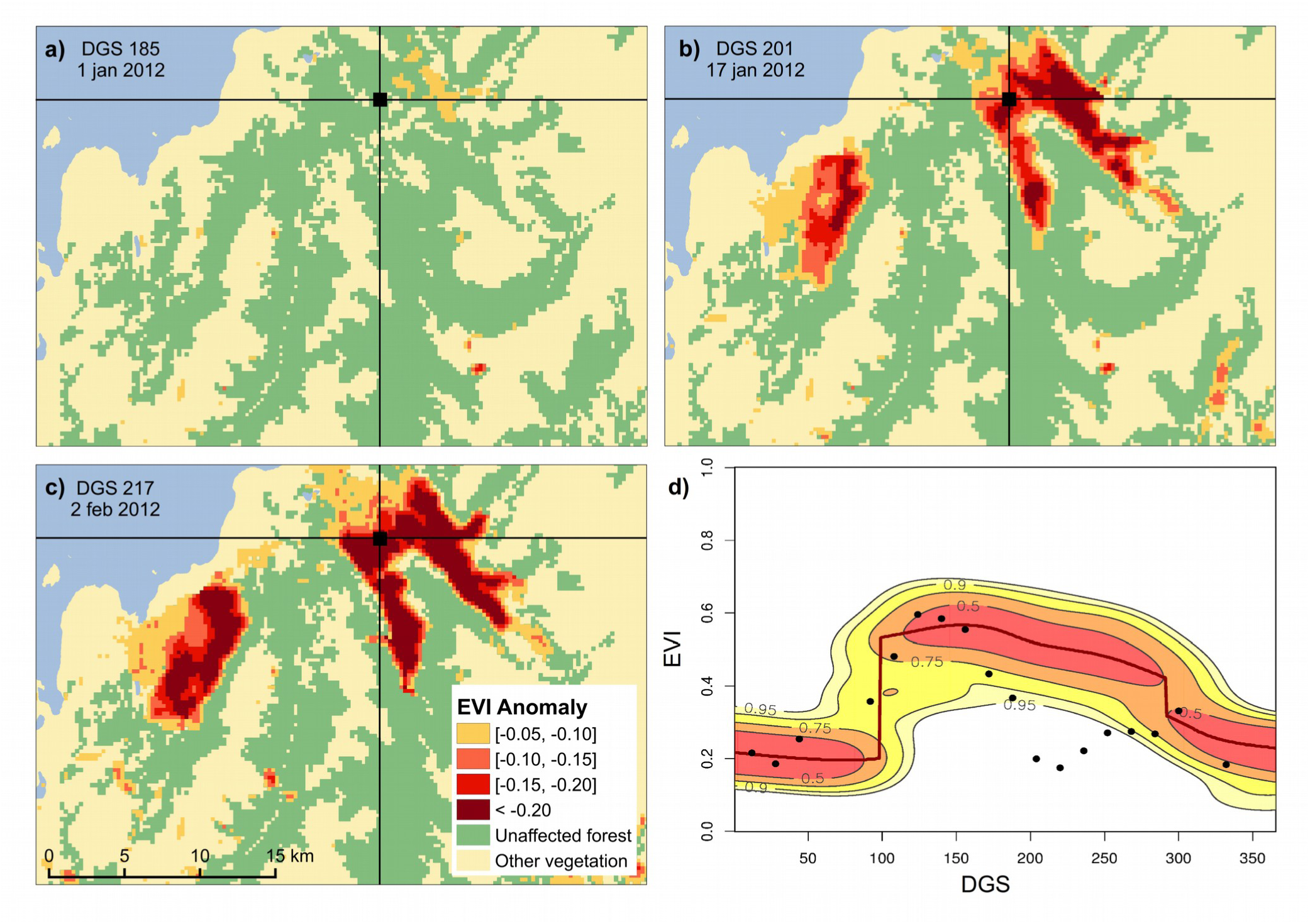
Example of npphen used for detecting and mapping foliage loss (“browning”) caused by an insect outbreak. Fig. 2a-c show EVI anomaly maps for different days of the growing season (DGS) displaying the spatial expansion and defoliation intensity of the *Ormiscodes amphimone* outbreak occurred during the growing season 2011-2012, which affected thousand of hectares of *Nothofagus pumilio* forests in Chilean Patagonia. Fig. 2d shows the density kernel plot and EVI anomalies for the growing season 2011-2012 at a single pixel located at 46.80° S and 72.49° W.

### Blooming desert

In this second example, we reconstructed the LSP cycle in the Atacama desert in Chile, and quantify the increase in plant biomass due to the “blooming desert” phenomenon. This phenomenon occurs when above-average rainfalls trigger a remarkable development and flowering of herbaceous plants. In this case, we used the GIMMS NDVI 3g product (Version 1), generated from the NOAA AVHRR satellites, consisting of bi-monthly NDVI records at 8 km pixel resolution and spanning from 1981 to 2015 (Pinzón and Tacker 2014). We downloaded 828 NDVI composites by using the R Package “gimms” (Detsch 2018), covering a large area of the Atacama Desert in Northern Chile. Maps showing positive anomalies or “greening” areas are shown in Fig. 3a-c. The LSP reconstruction for an specific pixel is shown in Fig. 3d, displaying the expected LSP of the desert: very low and flat throughout the growing season. This behavior was disrupted by unusual heavy rainfalls during year 2011, causing a massive bloom of herbaceous plants. Our algorithm detected and map these positive NDVI anomalies, allowing us to study the spatial and temporal spread of the blooming event.

**Figure 3.**
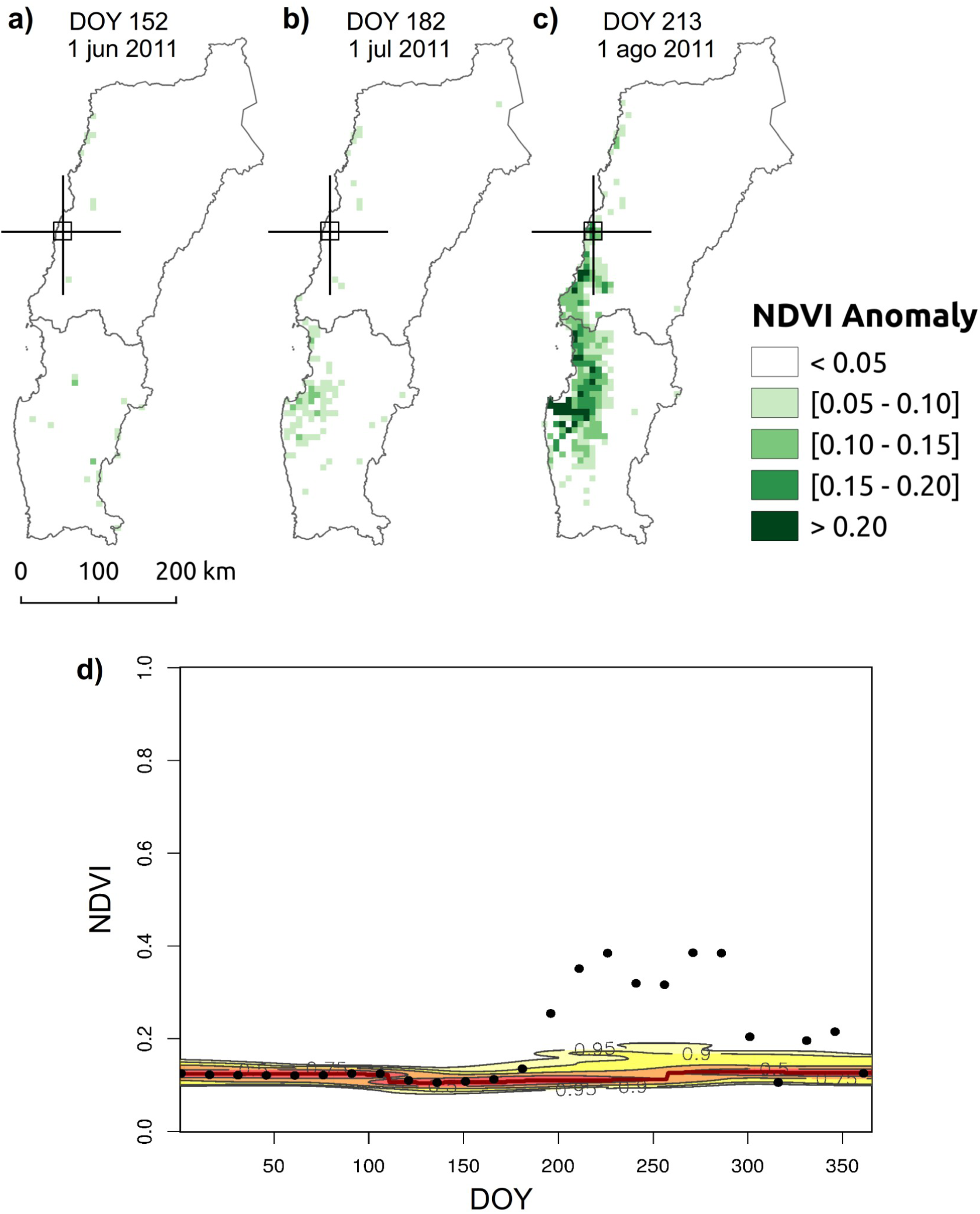
Example of npphen used for detecting and mapping a desert bloom (“greening”) caused by abnormal rainfall events occurred in 2012 in the hyper-arid Atacama Desert, Northern Chile. Fig. 3a-c show NDVI anomaly maps for different days of the year (DOY) displaying the spatial expansion and the development of a desert ephemeral grassland and flowers. Fig. 3d shows the the density kernel plot and anomalies for the year 2012 for a single pixel located at 27.96° S and 71.05° W.

## Conclusions and future directions

Our results show how the algorithms implemented in ‘npphen’ can successfully reconstruct the expected LSP for two contrasting ecosystems, using different data sources, and also detect and quantify anomalies caused by two different phenomena. Going back to the three challenges pointed out by Verbeselt et all. (2010), we will now explain how our algorithm overcome these issues. In the first place, ‘npphen’ takes advantage of long time series of satellite imagery to reconstruct the expected LSP for the complete time period used in training the algorithm. In this way we a) minimize the effect of particular disturbances by diluting their effects over data from multiple growing seasons, and b) increase the statistical power of the LSP estimation. Our approach has another relevant feature, which is the calculation of confidence intervals for the expected LSP. This is a key improvement because allows users to judge the magnitude of the anomaly on a probabilistic base, an aspect missing in current LSP reconstruction methods. de Beurs and Henebry (2010), after a thorough comparison of the different methods concluded that most of them have been developed for temperature and light limited vegetation, therefore non suitable for water-limited systems, and they all lack of a quality assessment of the significance or robustness of the model. ‘npphen’ overcomes both issues pointed out by this author.

In relation to the second challenge, our algorithm uses the KDE approach, and therefore, is completely independent of any particular functional form of the phenological curve. KDE methods are able to capture any functional form (Wand and Jones 1994), and therefore, can be applied at any type of vegetation. This has been illustrated by the two cases presented previously where ‘npphen’ was successfully applied to two contrasting vegetation types despite the high asymmetry in the distribution of the original data.

Finally, the LSP reconstructions using the complete time series and KDE methods ameliorate the data absence for some specific dates because these gaps can be partially filled with data from other seasons or contiguous dates. The case of *Nothofagus pumilio* forests is a good example of this situation. The study area in this case is commonly covered by clouds, and therefore, the time series has recurrent temporal gaps, which was not a limitation to perform the anomaly detection analysis. See Fig. 1b-c for an example of the estimation of LSP (Fig. 1c) from a pixel with multiple missing data (Fig. 1b).

In the near future we plan to improve the algorithm performance and include new functionalities. In relation to the former, despite the good performance of the current parallelization of some functions, we believe there is still room for reductions in the time consumed for some mapping functions. In relation to new functionalities, we are working on the implementation of simultaneous estimation of LSP anomalies for multiple growing seasons by means of a leave-one-out algorithm. We expect the ‘npphen’ packages will facilitate research on vegetation phenology and anomaly detection in natural systems, especially for unaccessible areas of the planet where remote sensing is the only viable alternative for monitoring.

## Acknowledgements

SAE is funded by CAPES-Conicyt FB-0002 (line 4), Fondecyt 1160370, and FIA PYT 2016-0203; ROCh was supported by CONICYT PAI N° 82140001 (2014) and Fondecyt Iniciación 11171046.

## Author’s contributions

SAE and ROCh conceived and designed research and prepared the manuscript. SAE and ROCh contributed with code to develop the ‘npphen’ package.

